# Nitric oxide regulates phagocytosis through S-nitrosylation of Rab5

**DOI:** 10.1101/2024.12.07.627306

**Authors:** Makoto Hagiwara, Hiroyuki Tada, Kenji Matsushita

## Abstract

Phagocytosis is mediated mainly by immune cells, such as macrophages, monocytes and neutrophils, that function to clear large pathogens including bacteria. The small GTP-binding protein Rab5 is crucial for clathrin-dependent endocytosis as well as phagocytosis. However, the role and mechanism of Rab5 activation during phagocytosis are poorly understood. Here we report that nitric oxide (NO), a novel regulator of Rab5, regulates phagocytosis through S-nitrosylation of Rab5. NO can promote phagocytosis by activating Rab5 in cultured cells, and it potently S-nitrosylates active Rab5 compared to inactive Rab5. Moreover, we demonstrate that two cysteine residues in the C terminus of Rab5 are S-nitrosylated and are important for phagocytosis. Experiments involving mice also showed that NO activates Rab5 and increases levels of S-nitrosylated Rab5 and that NO is involved in phagocytic bacterial clearance mediated by peritoneal macrophages. These data suggest that NO promotes S-nitrosylation of Rab5 to act as a novel Rab5 activator and a key regulator of phagocytosis.

**Highlights:**

1. NO promotes phagocytosis
2. NO directly activates Rab5 and increases Rab5 S-nitrosylation
3. S-nitrosylation C-terminal Rab5 cysteines residues is involved in phagocytosis
4. NO increases Rab5 S-nitrosylation and activity to promote bacterial clearance in vivo

## Introduction

Phagocytosis in mammals is carried out mainly by immune cells such as macrophages, monocytes and neutrophils, that function to clear large pathogens including bacteria, or large debris such as the remnants of dead cells or arterial fat deposits(1, 2). Phagocytosis is defined as the cellular engulfment of particles ≥0.5 μm in diameter and is a highly regulated process that involves cell-surface receptors (including cell adhesion proteins), intracellular signal transduction, actin remodeling and membrane trafficking(1, 2). For instance, binding of cellular receptors to bacteria triggers intracellular signaling, resulting in remodeling of the actin cytoskeleton and engulfment of bacteria by cell membranes that form the phagocytic cup. Subsequently, the leading edge of the growing cup closes, and a membrane vesicle (phagosome) develops(1, 2). Phagosomes containing bacteria undergo acidification and mature from Rab5-positive early to Rab7-positive late stages(2–5). Last, the phagosomes fuse with lysosomes to form phagolysosomes that degrade bacteria(2, 6, 7). However, many aspects of the mechanisms by which Rab5 regulates phagocytosis remain unclear.

Rab proteins are small GTPases that regulate vesicular transport in endocytosis and exocytosis(8–10). To date, over 60 distinct Rab proteins have been identified, and each protein is specifically associated with a particular organelle or pathway(8–10). Rab5, a member of the Rab family of GTPases, localizes to the plasma membrane and to endosomes(10–19). Several processes (e.g., budding, trafficking, tethering, and fusion of early endosomes) in eukaryotic cells require Rab5 for the coordination of endocytosis including phagocytosis that occurs via specific interactions involving over 30 effector proteins(2, 8–10, 20, 21). For example, Rabaptin-5(22, 23), EEA1(24), APPL1(25–27), Rabankyrin-5(28), and caveolin-1(29, 30) are known effectors of Rab5, and each protein is involved in regulation of endocytosis through interactions with Rab5. Cycling of Rab5 between an active (GTP-bound) and an inactive (GDP-bound) conformation is regulated by guanine nucleotide exchange factors (GEFs), GTPase-activating proteins (GAPs), and Rab GDP dissociation inhibitor (Rab GDI)(8–10). Rab5 replaces GDP with GTP through interactions with various GEF proteins such as Rabex-5(23), Rin1(31), Rin2(32), Rin3(33) and Alsin(34, 35) that recognize specific residues in the switch regions of Rab5 to facilitate GDP release. Conversion from the Rab5 GTP-bound form to the Rab5 GDP-bound form arises through GTP hydrolysis, which is controlled by the inherent GTPase activity of Rab5 but also by GAPs such as RabGAP-5(36),RN-tre(37), Armus (TBC-2)(38, 39), TBC1D17(40) and TBC1D18(41). Inactivated Rab5 separates from early endosome membranes (and also from the early phagosome membrane) and is retained in the cytosol by Rab GDI until the next round of the GTPase cycle begins(8–10). Post-translational modifications of Rab5 also affect its activity. Rab5 has long been known to undergo C-terminal geranylgeranylation that is thought to promote Rab5 membrane binding(9, 10, 42–45). Rab5 phosphorylation mediated by PKCε has been shown to be involved in regulating cell migration(46, 47). Mono-ubiquitination of Rab5 inhibits its effector binding and guanine nucleotide conversion(48). In this manner, post-translational modifications cause dynamic changes in Rab5 function.

Nitric oxide (NO) plays significant roles in various physiological processes including the regulation of immune responses(49), signal transduction(50–52), and blood pressure regulation(53). NO is produced from L-arginine by the NO synthase (NOS) isoforms neuronal NOS (nNOS, or NOS1), inducible NOS (iNOS, or NOS2), and endothelial NOS (eNOS or NOS3) in cells(51, 54). Recent studies reported that a NO moiety (NO•) to cysteines, a process called S-nitrosylation, represents a key signaling pathway that regulates protein functions(55–57). Protein S-nitrosylation is related to eNOS, nNOS, and iNOS expression levels in several tissues and is restricted to regions of the cell where NOS localizes (58–60). In addition, protein S-nitrosylation can be analyzed experimentally using NO donors such as S-nitrosoglutathione (GSNO), an agent that generates NO (61, 62). Many proteins, including caspase-3(63), MyD88(64), beta-actin(65), caveolin-1(66), and PTEN(67, 68), are reported to be S-nitrosylated. We previously reported that S-nitrosylation of NSF, an ATPase that is essential for activating membrane fusion machinery, regulates exocytosis of Weibel-Palade bodies and platelet granule (69, 70). However, the impact of NO on regulatory factors that control endocytosis and phagocytosis remains unclear. In this study, we describe a novel molecular mechanism by which NO regulates phagocytosis through S-nitrosylation of Rab5.

## Materials and Methods

### Cell culture

RAW264 cells (RIKEN BioResource Research Centre, Ibaraki, Japan, Cell No. RCB0535) were cultured in Dulbecco’s modified Eagle’s medium (DMEM) supplemented with 10% fetal bovine serum (FBS), penicillin (100 IU/ml) and streptomycin (100 IU/ml) at 37 °C in a humidified atmosphere with 5% CO_2_. HEK293T cells (Japanese Collection of Research Bioresources Cell Bank, Ibaraki, Japan, Cell No. JCRB9068) were cultured in Dulbecco’s modified Eagle’s medium (DMEM) supplemented with 10% fetal bovine serum (FBS), penicillin (100 IU/ml) and streptomycin (100 IU/ml) at 37 °C in 5% CO_2_.

### Vector constructs

iNOS cDNA was subcloned into the pCI vector. GFP-Rab5 in pcDNA3 and GST-Rab5Q79L, GST-Rab5 (wild) and GSTRab5S34N pGEX-2T constructs were kindly provided by Dr. Y. Yamamoto (Tokyo University of Agriculture, Tokyo, Japan). Rab5 was cloned into the pet30a vector to obtain His-Rab5. GST-Rab5C19A, GST-Rab5C63A, GST-Rab5C212A, GST-Rab5C213A and GST-Rab5C212A/C213A in pGEX-2T were constructed by mutating GST-Rab5 (wild type) in pGEX-2T using a Quik-Change Site-Directed Mutagenesis Kit (Agilent Technologies, San Diego, USA). The mutated Rab5 genes were then cloned into the pCMV-HA vector. The GST-R5BD vector was kindly provided by Dr. G. Li (University of Oklahoma Health Science Center, Oklahoma City, USA)(71).

### Antibodies

Antibodies were obtained from the following sources: anti-mouse HA and anti-rabbit HA (Sigma-Aldrich, St. Louis, MO USA); anti-rabbit IgG-Alexa 555 and anti-rabbit IgG-Alexa 633 (Life Technologies, Carlsbad, CA USA); anti-mouse Rab5, anti-rabbit Rab5, anti-mouse GFP, and anti-rabbit GFP (Novus, Littleton, CO USA); anti-mouse IgG-HRP and anti-rabbit IgG-HRP (IBL, Gunma, Japan); anti-GST HRP conjugate (GE Healthcare, Fairfield, CT USA); anti-mouse GAPDH (MBL, Nagoya, Japan); anti-mouse His (SinoBiological, North Wales, PA USA); and anti-rabbit iNOS (NOS2) (Santa Cruz Biotechnology).

### Transfection

RAW264 cells were transfected with plasmids using Lipofectamine 3000 (Life Technologies, Carlsbad, CA USA) and incubated for 3 h. The medium was then aspirated and replaced with fresh medium. The cells were once again transfected with plasmids using Lipofectamine 3000.

### Phagocytosis assay

Transfected cells or cells treated with S-nitrosoglutathione (GSNO) or NG-nitro-L-arginine methyl ester hydrochloride (L-NAME HCL, hereafter abbreviated as L-NAME) in 96-well plates were pre-incubated with serum-free DMEM without phenol red for 1 h at 37 °C. The medium was then aspirated and 1 mg/mL pHrodo red-labeled *S. aureus* BioParticles (Life Technologies, Carlsbad, CA USA), a phagocytosis marker that was diluted with serum-free DMEM without phenol red, was added to the cells. The cells were incubated for the indicated times at 37 °C, and phagocytic uptake was analyzed by measuring fluorescence signal intensity with a SpectraMax m3 multimode microplate reader (Molecular Devices Japan, Tokyo, Japan) (excitation wavelength: 560 nm / fluorescence wavelength: 585 nm). Fluorescence microscopy observations were made under the same conditions.

### SDS-PAGE and western blotting

SDS-PAGE and western blotting were performed using experimental conditions reported by Hagiwara(72, 73), except that a E-T520L e-PAGEL (5-20% gradient SDS-polyacrylamide gel, catalog number 2331830, ATTO Corporation) was used.

### Bacterial protein expression and purification

Bacterial expression and purification of GST-R5BD was performed as previously described(12, 15, 30). Purified GST-R5BD was stored at −80 °C until use.

Purified His-Rab5 was prepared using a standard procedure. In brief, the Rab5 in the pet30a vector was transformed into *E. coli* BL21-CodonPlus (DE3) RIL competent cells (Agilent Technologies, San Diego, CA USA) and protein expression was induced with 1 mM isopropyl thio-β-D-galactoside (IPTG) for 3 h at 37 °C. His-Rab5 was purified from bacterial cell extracts by passage over a His Trap HP column (GE Healthcare, Fairfield, CT USA) and elution with imidazole according to the manufacturer’s instructions. Desalting and buffer exchange into PBS was conducted with an Amicon Ultra device (Merck, Darmstadt, Germany). Purified His-Rab5 was stored at −80 °C until use.

### Immunoprecipitation

Cells were lysed on ice in buffer (10 mM Tris, pH 7.6, 150 mM NaCl, 5 mM MgCl_2_, 1% Triton X-100) containing protease inhibitors and clarified by centrifugation. Cell lysates were incubated with an antibody against the indicated target protein with rotation for 1 h at 4 °C. Protein A-agarose beads were then added to the cell lysates and the mixtures were incubated for an additional 1 h at 4 °C. After washing, the immunoprecipitated proteins were dissolved in SDS-sample buffer and analyzed by western blotting.

### Immunostaining

After washing with PBS, cells were fixed in 4% paraformaldehyde for 10 min, permeabilized with 0.1% Triton X-100 for 10 min, and blocked with blocking reagent N101 (NOF corporation, Tokyo, Japan) for 1h at room temperature. The cells were then incubated with primary antibodies against the indicated target proteins, followed by incubation with the respective fluorescent secondary antibodies. After extensive washing, cells were mounted on coverslips using Shandon Immu-Mount (Thermo, Tokyo, Japan). Cells were observed with confocal fluorescence microscopy (Carl Zeiss, Jena, Germany) and images were analyzed using ZEN 2009 imaging software.

### Detection of S-nitrosylation

A biotin-switch assay was carried out using a S-Nitrosylated Protein Detection Kit (Cayman, Michigan, USA) according to the manufacturer’s instructions. Biotinylated proteins were precipitated with streptavidin agarose beads and western blotting was performed to detect SNO-Rab5.

### GST-R5BD pull down assay

A GST-R5BD pull-down assay was performed as previously described(12, 15, 30). Briefly, glutathione Sepharose beads were coated with 30 µg of GST-R5BD. The beads were then incubated with lysates of freshly-transfected HEK293T cells or purified His-Rab5 for 30 min at 4 °C. The beads were washed and subjected to western blotting.

### GDP release assay and GTP-binding assay

A GDP release assay using mant-GDP and a GTP-binding assay using mant-GTP were performed as previously reported(74–76). Fluorescence measurements were made using a SpectraMax m3 multimode microplate reader (excitation wavelength: 355 nm / fluorescence wavelength: 448 nm).

### Subcellular fractionation

Cells expressing each of the HA-Rab5 mutants were homogenized in HEPES buffer (20 mM HEPES (pH 7.4), 100 mM NaCl, 5 mM MgCl_2_) containing protease inhibitors. The homogenates were centrifuged at 800×g for 10 min at 4 °C to remove cell debris. The supernatant was further centrifuged at 200,000×g for 30 min, yielding supernatant (S) and pellet (P) fractions. P was subsequently resuspended in HEPES buffer. S and P fractions were analyzed by western blotting.

### Visualization of S-nitrosylated proteins

A biotin-switch assay was performed using a S-Nitrosylated Protein Detection Kit (Cayman, Michigan, USA) according to the manufacturer’s instructions. Cells were observed with confocal fluorescence microscopy and images were analyzed using ZEN 2009 imaging software.

### In vivo Rab5 activity measurement and Rab5 S-nitrosylation analysis

Female mice (C57BL/6J JmsSlc) were purchased from Japan SLC, Inc. (Shizuoka, Japan). To detect Rab4 S-nitrosylation, mice were injected intravenously with lipopolysaccharide (LPS) or LPS and L-NAME, and sacrificed 24 hours later. The spleens were immediately collected and homogenized for analysis of Rab5 S-nitrosylation using a S-Nitrosylated Protein Detection Kit (Cayman Chemical, Ann Arbor, MI USA) according to the manufacturer’s instructions and for measurement of Rab5 activity using a GST-R5BD pull down assay as described above.

The protocols for all experiments involving animals were approved by the Animal Experiment Committee of the National Center for Geriatrics and Gerontology (Approval number 26-21-R1) and the experiments were conducted in accordance with the animal experiment regulations of the National Center for Geriatrics and Gerontology.

### Staphylococcus aureus culture conditions

*S. aureus* ATCC 35896 was cultured aerobically in brain heart infusion (BHI) broth at 37 °C for 12 h. Bacterial cultures in the log phase of growth were then centrifuged for 15 min at 8500 × *g*, washed three times with PBS, and cultured aerobically on BHI agar plates at 37 °C for 12 h. The number of colony forming units (CFU)/mL was then determined.

### Clearance of *S. aureus* by intraperitoneal macrophage transfer

Donor female C57BL/6 mice were intraperitoneally injected with 4% thioglycolate (2 mL/mouse). Peritoneal macrophages collected 4 days later were cultured in 6-well plates with DMEM/F-12 containing 10% FBS with 4 × 10^6^ cells/well. Macrophages were treated with 1 mM L-NAME for 24 h, and washed three times with PBS. Adherent cells were collected using Accutase (Nacalai Tesque, Kyoto, Japan). Host female C57BL/6 mice were intraperitoneally injected with *S. aureus* (3 × 10^9^ CFU/mice) together with the macrophages (4 × 10^6^ CFU/mice). After 6 h, *S. aureus* in the peritoneal lavage fluid was cultured on BHI agar and the number of CFU was determined. These experiments were approved by the Animal Experiment Committee of Tohoku University (Approval number 2014-037) and were conducted in accordance with the animal experiment regulations of Tohoku University.

### Statistical analysis

Statistical analysis of results from experiments having two samples was conducted using an F-test followed by a t-test. Statistical analysis of results from experiments having more than three samples was conducted using the Holm–Bonferroni method. p-values ≤0.05 were considered significant.

## Results

### NO regulates phagocytosis

We first examined the effect of nitric oxide (NO) on phagocytosis in macrophage-like RAW264 cells. GSNO is a source of NO (NO donar) and an important mediator of NO signaling. It can generate NO through processes such as heat, photodecomposition, metal ions, reducing agents, and other mechanisms. From this, we investigated the effect of NO on phagocytosis using GSNO. RAW264 cells were preincubated with GSNO or GSH (control) for 1h and assayed for uptake of pHrodo Red *S. aureus* BioParticles, a marker of phagocytosis. RAW264 cells incubated with GSNO had increased phagocytic uptake compared to those incubated with GSH (Fig. 1A and B). Phagocytosis was also upregulated in RAW264 cells with overexpression of iNOS (inducible nitric oxide synthase; Fig. 1C and D). Stimulation with LPS is known to dramatically increase iNOS expression and NO production in cells(77, 78). To clarify whether LPS-stimulated NO production promotes phagocytosis, we conducted experiments using L-NAME, a nonselective NOS inhibitor. The results showed that stimulation with LPS alone for 24h promoted phagocytosis, whereas phagocytosis was decreased in RAW264 cells treated with L-NAME alone or with both LPS and L-NAME for 24h (Fig. 1E and F). Taken together, these results suggest that NO upregulates phagocytosis in RAW264 cells.

**Fig. 1.**
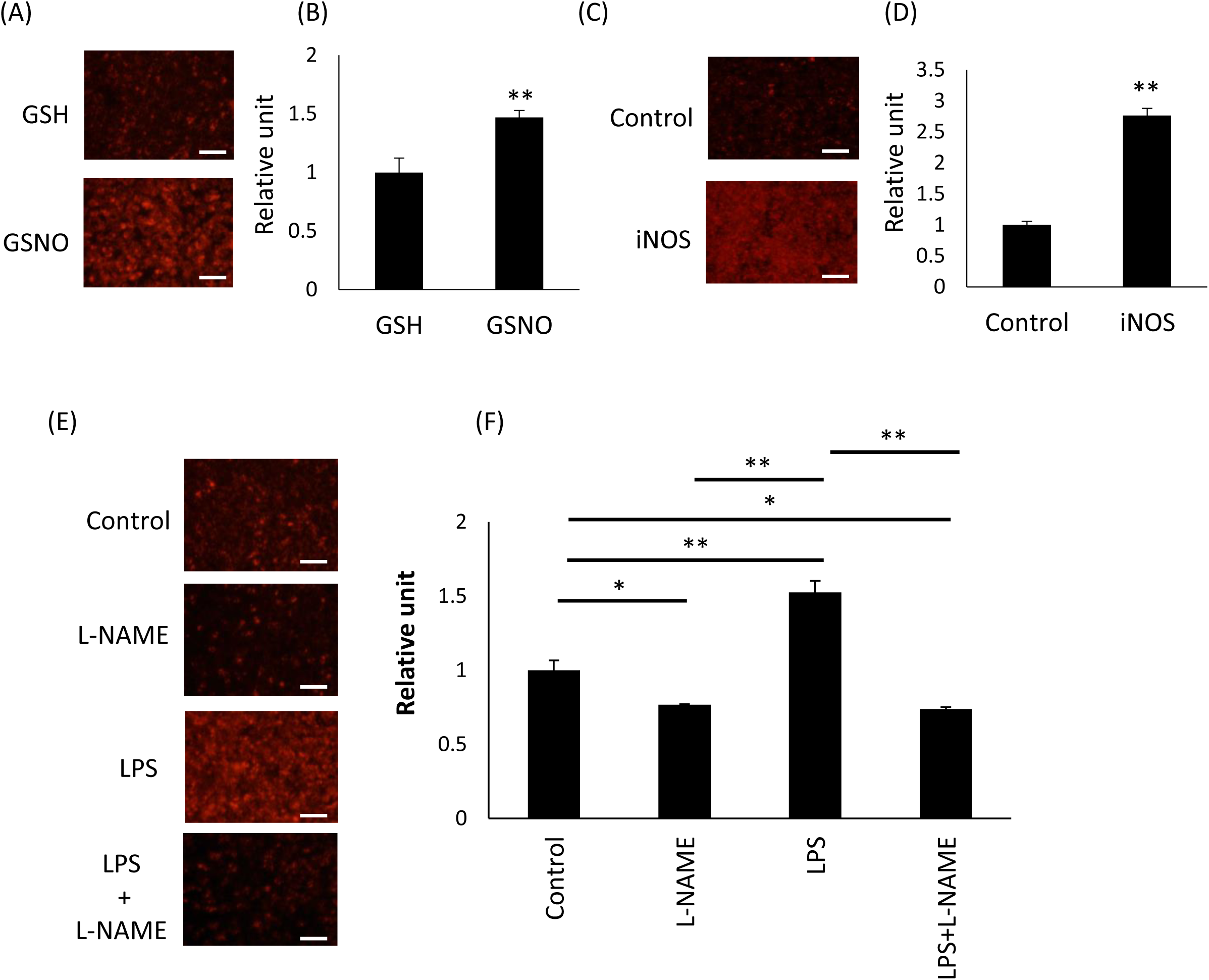
NO regulates phagocytosis. **(A and B)** RAW264 cells were incubated with 100 μM GSNO, a NO donor, or glutathione (GSH) for 1 h. The cells were then incubated with pHrodo Red *S. aureus* BioParticle conjugates for phagocytosis for 1h. (A) Images were taken with a fluorescence microscope. Scale bar: 200 µm. (B) Values measured with a fluorescent microplate reader. Each value in the graph is the mean ± SD of three independent experiments. **p < 0.01. (C and D) RAW264 cells transfected with an iNOS vector or a control vector were incubated for 1h 37 °C with pHrodo Red *S. aureus* BioParticle conjugates for phagocytosis. (C) Images taken with a fluorescence microscope. Scale bar: 200 µm. (D) Values measured with a fluorescent microplate reader. Each value in the graph is the mean ± SD of three independent experiments. **p < 0.01. (E and F) RAW264 cells were incubated with 1 mM L-NAME, 100ng/ml LPS, 100ng/ml+L-NAME, or dimethyl sulfoxide (DMSO) alone (control) for 24 h. The cells were then incubated with pHrodo Red *S. aureus* BioParticle conjugates for phagocytosis. (E) Images taken with a fluorescence microscope. Scale bar: 200 µm. (F) Values measured with a fluorescent microplate reader. Each value in the graph is the mean ± SD of three independent experiments. *p < 0.05, **p < 0.01.

### iNOS interacts with Rab5

Next, we examined whether iNOS interacts with Rab5. RAW264 cells transfected with HA-Rab5 or control vector were incubated with or without 100 ng/ml LPS for 16 h. iNOS was co-immunoprecipitated with HA-Rab5 in lysates from LPS-stimulated RAW264 cells (Fig. 2A). To investigate whether iNOS interacts with inactive and/or active Rab5, we performed a GST pull-down assay of LPS-stimulated RAW264 cells expressing either Rab5 (Q79L), an active mutant, or Rab5 (S34N), an inactive mutant. The results revealed that iNOS binds more strongly to active Rab5 (Q79L) than to inactive Rab5 (S34N) (Fig. 2B). Next, we observed the intracellular localization of iNOS and Rab5 in RAW264 cells with confocal fluorescence microscopy. In RAW264 cells transfected with iNOS and HA-Rab5, iNOS colocalized with HA-Rab5 (Fig. 2C). These data suggest that iNOS interacts with Rab5.

**Fig. 2.**
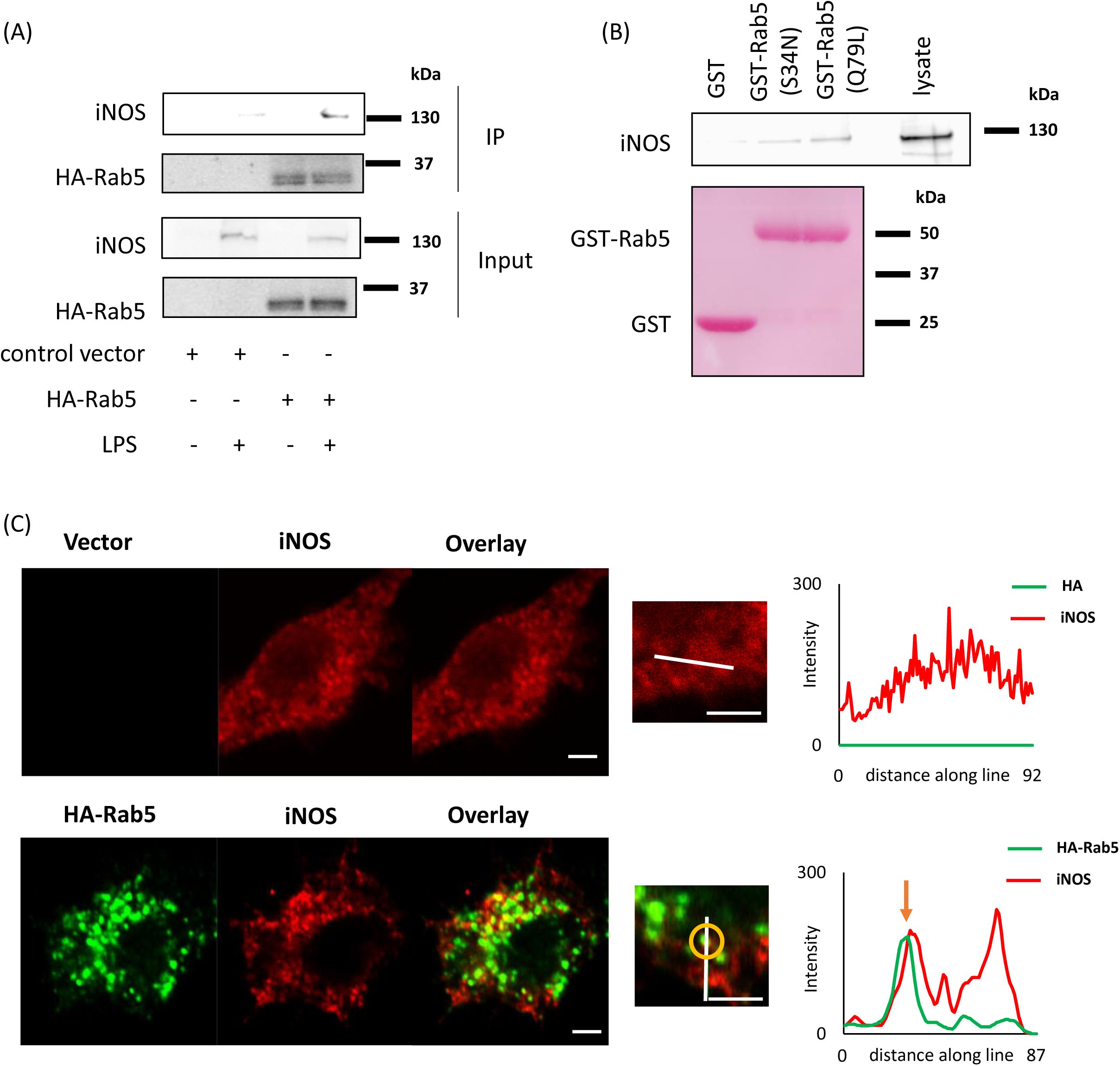
iNOS interacts with Rab5. (A) RAW264 cells transfected with HA-Rab5 or control vector were incubated with 100 ng/ml LPS for 16 h. HA-Rab5 in lysates from LPS-stimulated RAW264 cell lysates was then immunoprecipitated and iNOS binding to Rab5 was assessed with western blotting. (B) A pull-down assay with LPS-stimulated RAW264 cell lysates was performed using GST, GST-Rab5(S34N), and GST-Rab5(Q79L). Binding of iNOS to the beads was assayed with western blotting. GST, GST-Rab5(S34N), and GST-Rab5(Q79L) on the PVDF membrane were stained with Ponceau S. (C) RAW264 cells transfected with HA-Rab5 (wild type) and iNOS were fixed and immunostained with anti-HA and anti-iNOS antibodies. The cells were then observed with confocal fluorescence microscopy. Scale bar: 5 µm. Fluorescence intensities of HA-Rab5 (green) and iNOS (red) measured along the line in the left panel.

### NO directly increases Rab5 activity

Since NO activated phagocytosis (Fig. 1A-F), we hypothesized that NO would also promote Rab5 activity. We used a Rabaptin-5 Rab5 binding domain-based GST pull down assay (GST-R5BD pull down assay) to evaluate Rab5 activation(71). We found that Rab5 activity in HEK293 cells dose- and time-dependently increased in response to GSNO (Fig. 3A and B). Moreover, HEK293 cells expressing iNOS and HA-Rab5 exhibited increased Rab5 activity (Fig. 3C). To further examine whether NO directly enhanced Rab5 activity, we used purified recombinant His-Rab5. Again we found that GSNO dose- and time-dependently increased activity of purified Rab5 (Fig. 4A and B). These data suggested that NO indeed regulates Rab5 activity.

**Fig. 3.**
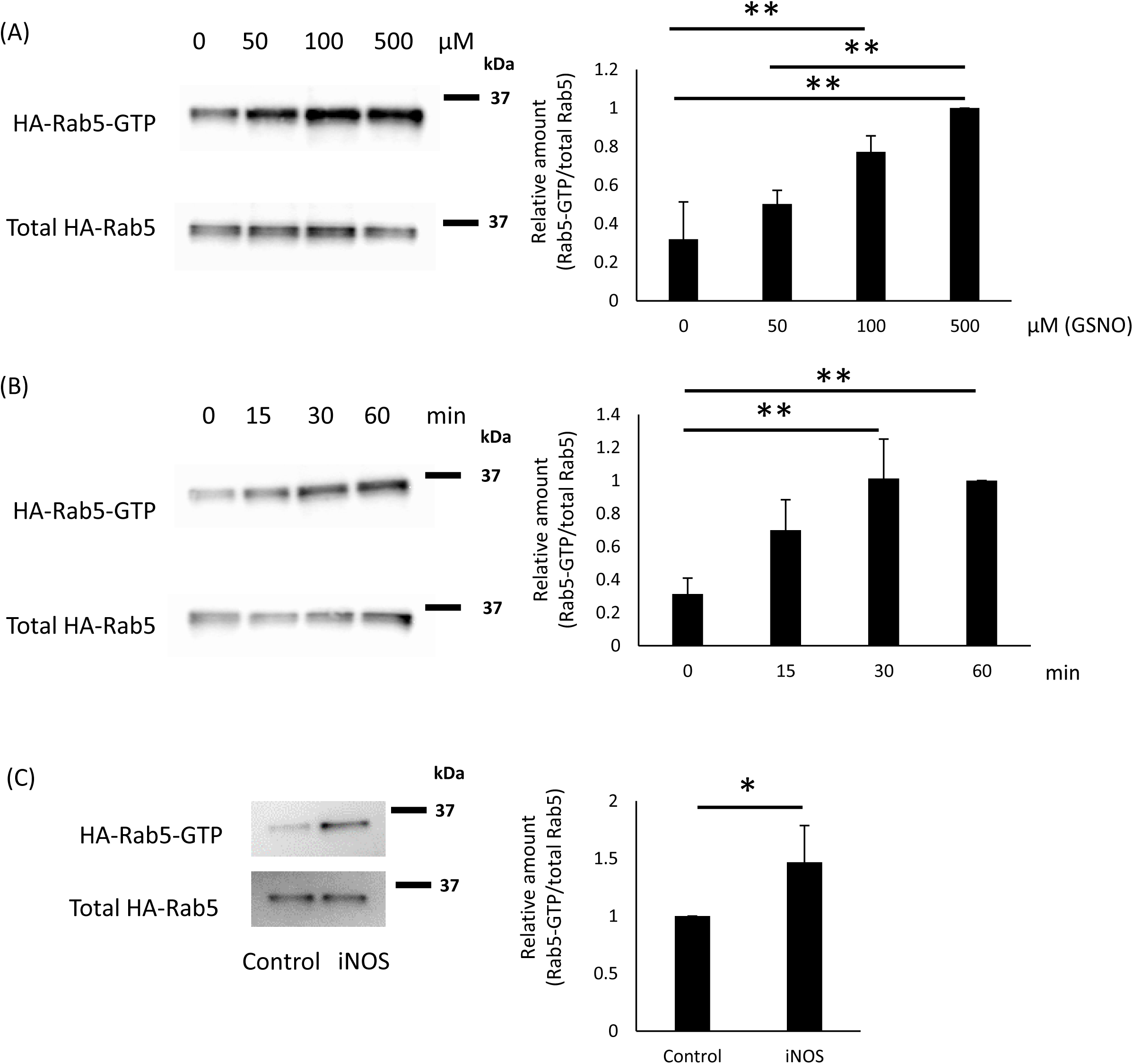
NO increases Rab5 activity. (A) HEK293 cells transfected with an HA-Rab5 vector were incubated with 0-500 μM GSNO for 60 min. Rab5-GTP was assessed with a GST-R5BD pull-down assay. Each value in the graph is the mean ± SD of four independent experiments. **p < 0.01. (B) HEK293 cells transfected with an HA-Rab5 vector were incubated with 100 μM GSNO for 0-60 min. Rab5-GTP was assessed with a GST-R5BD pull-down assay. Each value in the graph is the mean ± SD of three independent experiments. **p < 0.01. (C) HEK293 cells were transfected with iNOS and HA-Rab5 vectors and Rab5-GTP was assessed with a GST-R5BD pull-down assay. Each value in the graph is the mean ± SD of four independent experiments. *p < 0.05.

**Fig. 4.**
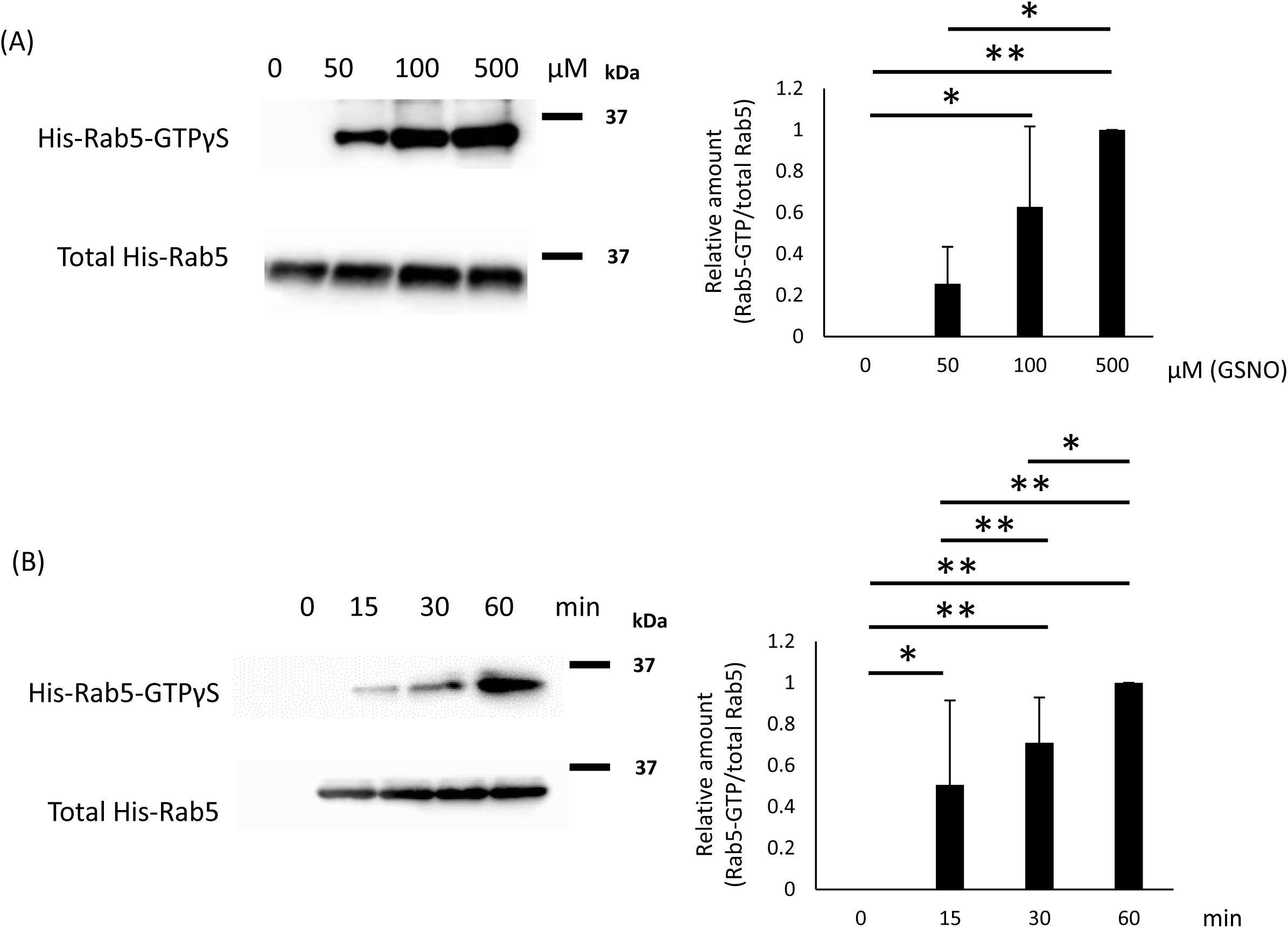
NO directly increases Rab5 activity. (A) Purified recombinant His-Rab5 (Wild) was incubated with 0-500 μM GSNO and GTPγS for 30 min at room temperature and Rab5-GTPγS was assessed with a GST-R5BD pull-down assay. Each value in the graph is the mean ± SD of three independent experiments. *p < 0.05, **p < 0.01. (B) Purified recombinant His-Rab5 (Wild) was incubated with 100 μM GSNO and GTPγS for 0-60 min at room temperature, and Rab5-GTPγS was assessed with a GST-R5BD pull-down assay. Each value in the graph is the mean ± SD of three independent experiments. *p < 0.05, **p < 0.01.

We next analyzed the Rab5-mediated GDP/GTP nucleotide exchange reaction in response to NO. Aqueous solutions of GSNO are pink in color, which interferes with fluorescence measurements taken with a microplate reader. Therefore, in the next experiment, DEA-NONOate, a colorless NO donor with a short half-life that rapidly releases nitric oxide, was used. Mant-GDP (fluorescent labeled GDP analog)-loaded recombinant His-Rab5 was incubated with DEA-NONOate. DEA-NONOate treatment promoted GDP dissociation from Rab5 (Fig. 5A). Meanwhile, incubation of Rab5 with mant-GTP, a fluorescent GTP nucleotide analog, and DEA-NONOate resulted in binding of GTP to Rab5 (Fig. 5B). These results indicate that NO is directly involved in the GDP/GTP exchange reaction of Rab5.

**Fig. 5.**
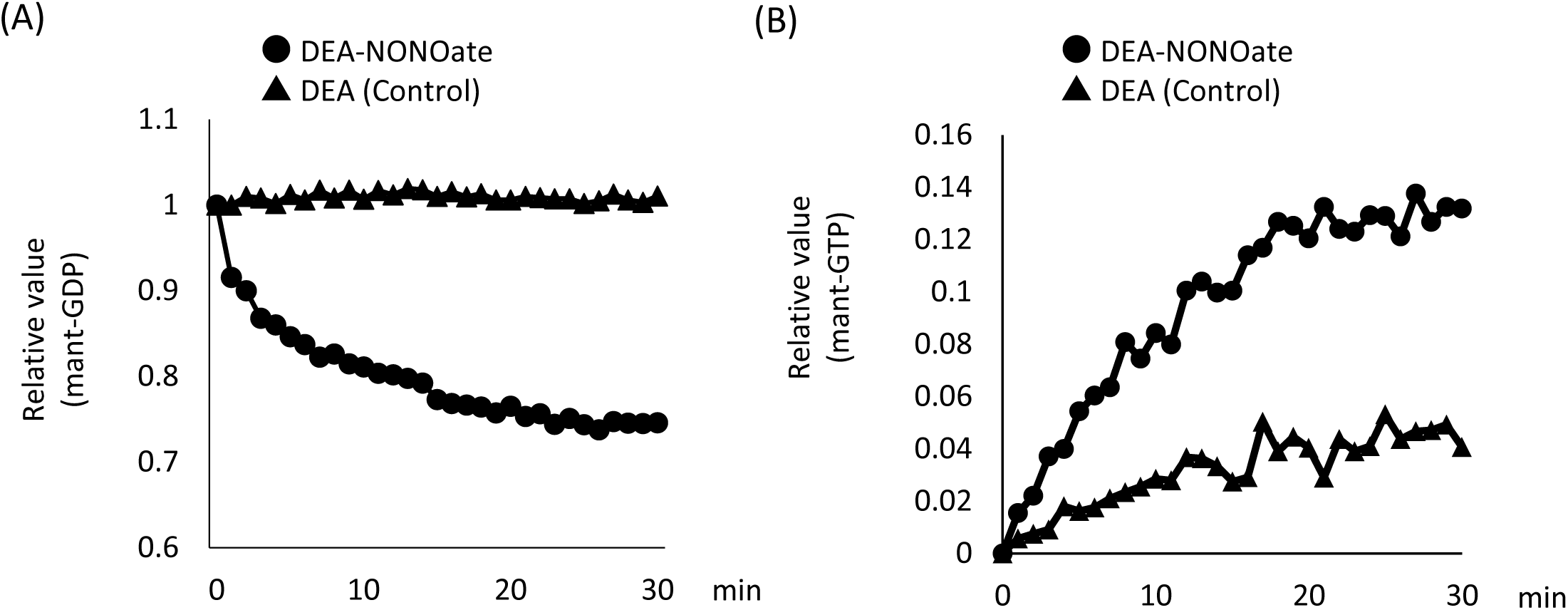
NO is a guanine-nucleotide exchange factor for Rab5. (A) Mant-GDP-loaded recombinant His-Rab5 was incubated with DEA-NONOate. The fluorescence emission was monitored for 30 min. (B) Recombinant His-Rab5 was incubated with DEA-NONOate and mant-GTP. The fluorescence emission was monitored for 30 min.

### NO S-nitrosylates Rab5

The biotin switch assay is a powerful tool for detecting S-nitrosylated proteins(79). We next measured S-nitrosylation of Rab5. HEK293 cells transfected with an HA-Rab5 vector were incubated with GSNO and Rab5 S-nitrosylation was assessed using a biotin-switch assay. Increased S-nitrosylation of Rab5 expressed in HEK293 cells occurred dose- and time-dependently following GSNO treatment (Fig. 6A and B). To investigate whether S-nitrosylation is related to the activity status of Rab5, we detected S-nitrosylation of active Rab 5 and inactive Rab5 mutants. Purified GST-Rab5 (Q79L) and GST-Rab5 (S34N) were incubated with GSNO and S-nitrosylation of Rab5 was assessed using a biotin-switch assay. Both active Rab5 (Q79L) and inactive Rab5 (S34N) were S-nitrosylated, but the levels of S-nitrosylation were higher for active Rab5 (Q79L) than for inactive Rab5 (S34N) (Fig. 6C). In addition, HA-Rab5 showed increased S-nitrosylation in iNOS- and HA-Rab5-expressing HEK293 cells (Fig. 6D). These data suggested that Rab5 is S-nitrosylated by NO.

**Fig. 6.**
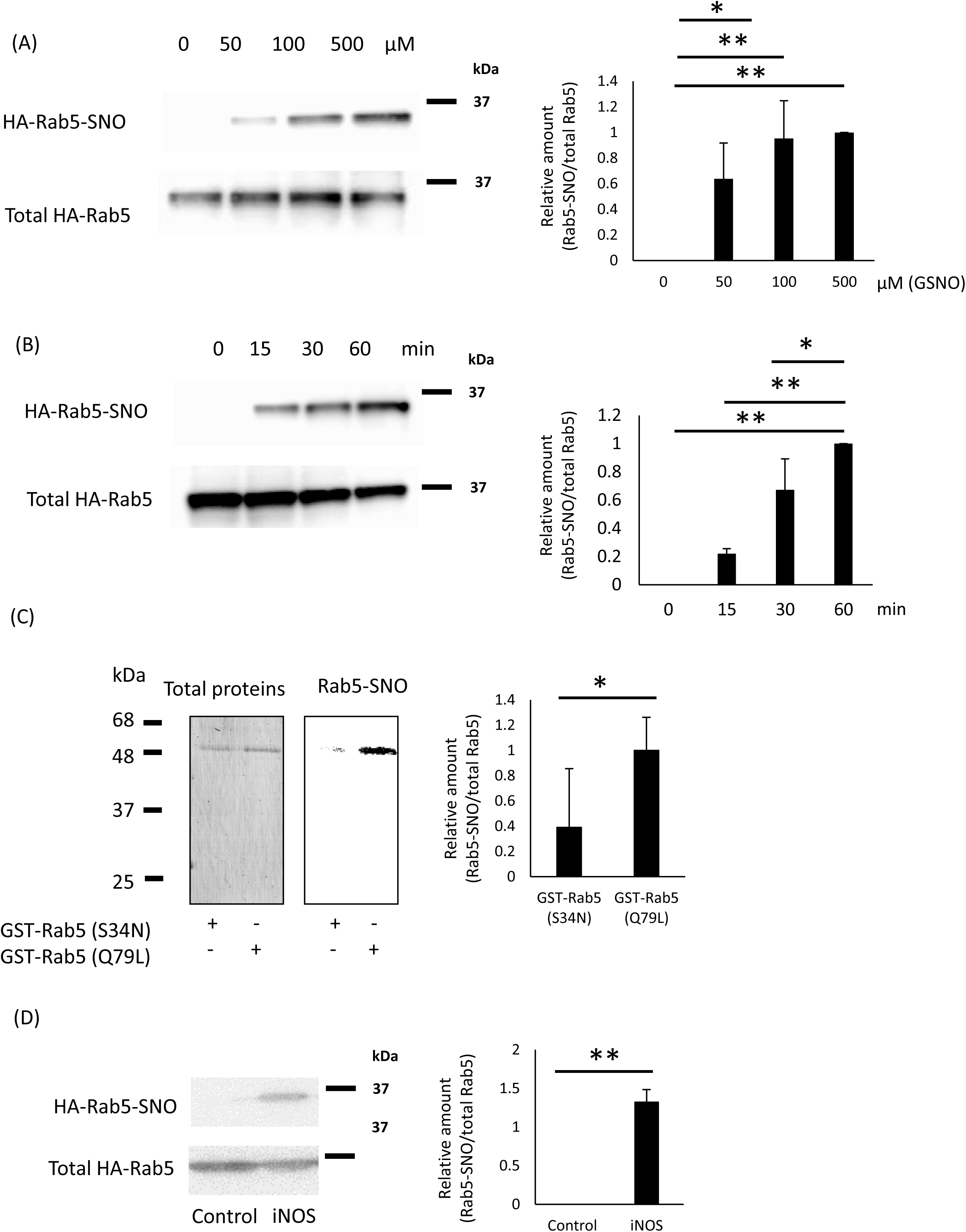
Rab5 is nitrosylated. (A) HEK293 cells transfected with an HA-Rab5 vector were incubated with 0-500 μM GSNO for 60 min. S-nitrosylation of Rab5 was assessed using a biotin-switch method. Each value in the graph is the mean ± SD of three independent experiments. *p < 0.05, **p < 0.01. (B) HEK293 cells transfected with an HA-Rab5 vector were incubated with 100 μM GSNO for 0–60 min. S-nitrosylation of Rab5 was assessed using a biotin-switch method. Each value in the graph is the mean ± SD of three independent experiments. *p < 0.05, **p < 0.01. (C) Purified recombinant GST-Rab5(Q79L) or GST-Rab5(S34N) were incubated with 500 μM GSNO at room temperature for 60 min, and S-nitrosylation was assessed with a biotin-switch method. Each value in the graph is the mean ± SD of three independent experiments. *p < 0.05. (D) HEK293 cells were transfected with iNOS and HA-Rab5 vectors and the S-nitrosylation of Rab5 was assessed using a biotin-switch method. Each value in the graph is the mean ± SD of five independent experiments. **p < 0.01.

### S-nitrosylation of Rab5 cysteine residues

After demonstrating that Rab5 is S-nitrosylated (Fig. 6A-D), we next identified the S-nitrosylation sites. S-nitrosylation has consistently been reported to occur at cysteine residues(55, 80, 81). The mouse Rab5 amino acid sequence has four cysteine residues: C19, C63, C212, and C213. We mutated these cysteine residues to alanine, which does not undergo S-nitrosylation, and also created mutants in which the two cysteine residues on the C-terminal side were mutated to alanine. (Fig. 7A). To analyze the subcellular distribution of each HA-Rab5 mutant, the cytosolic (S) and membrane-containing particulate (P) fractions were prepared by ultracentrifugation of cell lysates. All HA-Rab5 variants, including HA-Rab5C212A/C213A (double mutation with two C-terminal cysteines mutated to alanine Rab5), were detected in both the S and P fractions (Fig. 7B). This result was unexpected given that previous reports showed that Rab5 binds membranes via lipid modifications at the cysteines in the C-terminal of Rab5 (see Discussion).

**Fig. 7.**
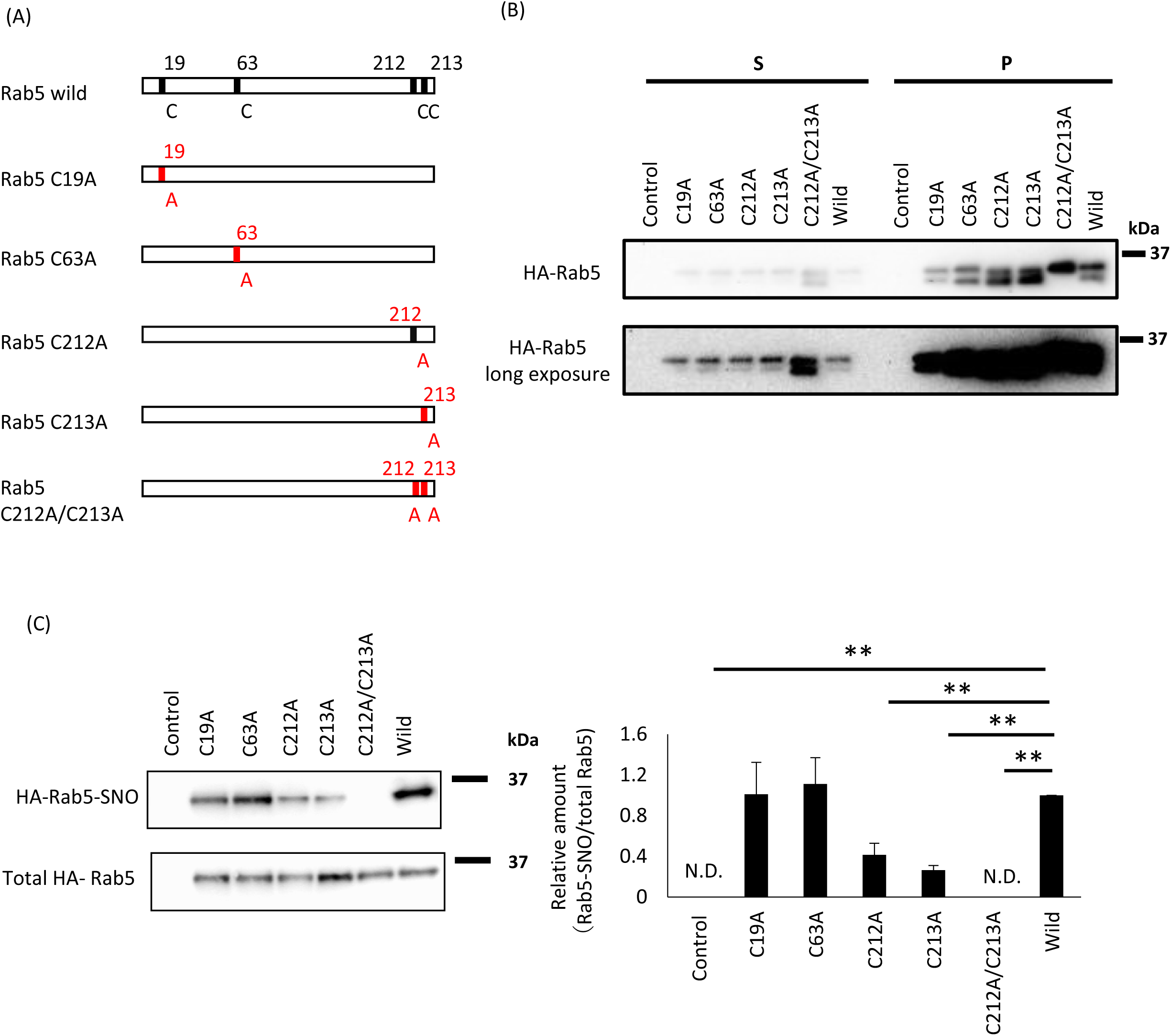
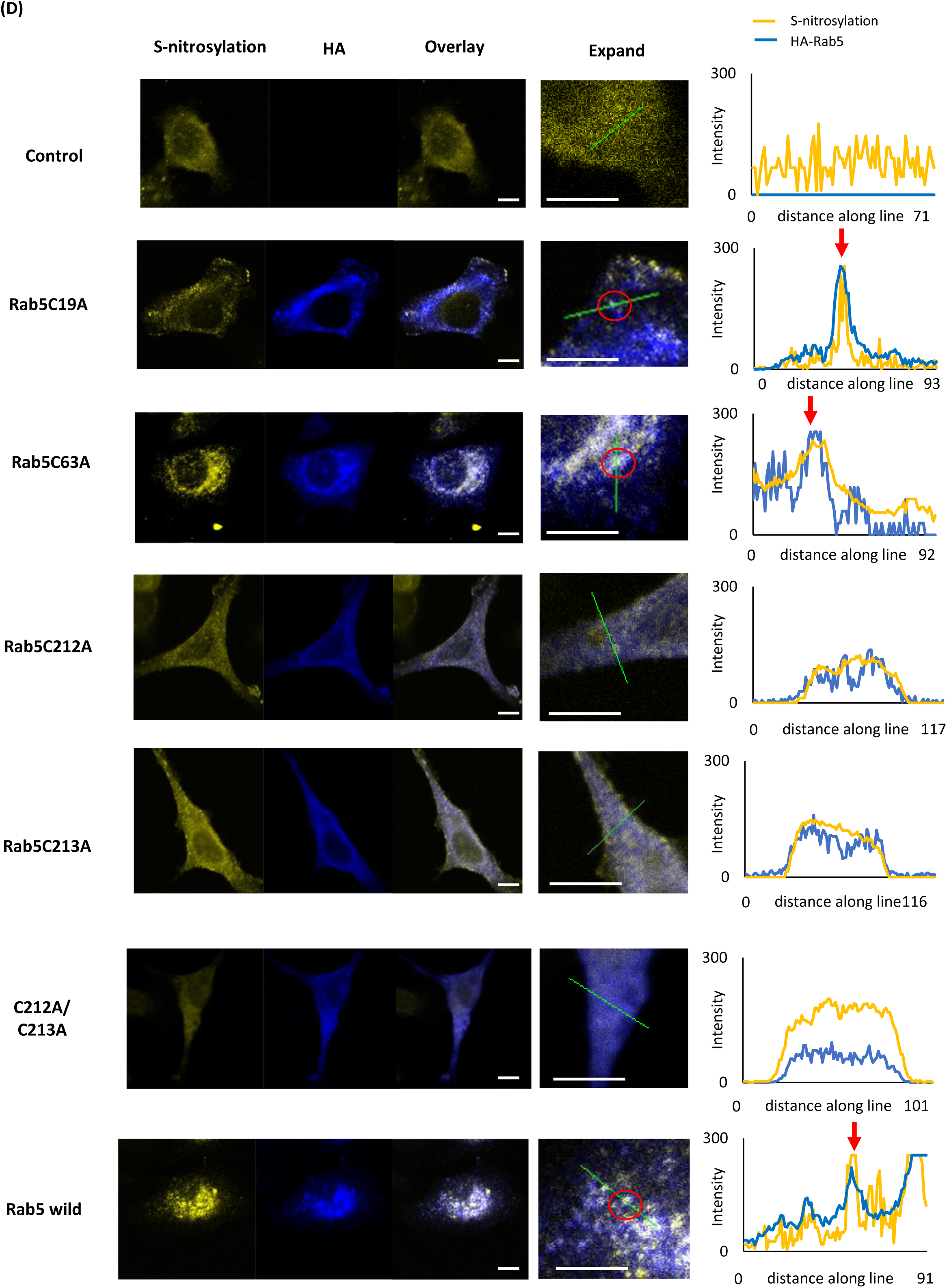
Rab5 cysteine residues C212 and C213 are S-nitrosylated. (A) Schematic of Rab5 mutants. (B) HEK293 cells transfected with the indicated HA-Rab5 mutant vector, HA-Rab5 wild type vector or control vector were homogenized and the cell lysates were ultracentrifuged. The resulting supernatant (S) and pellet (P) fractions were analyzed by western blotting. (C) HEK293 cells transfected with the indicated HA-Rab5 mutant vector, HA-Rab5 wild type vector or control vector were incubated with 100 μM GSNO for 1 h. S-nitrosylation of HA-Rab5 was detected by a biotin-switch assay. Rab5-SNO levels were normalized to total HA-Rab5 levels and quantified using ImageJ. The graph shows the mean ± SE of three independent experiments, **p < 0.01 versus HA-Rab5 wild-expressing cells; N.D., not detected. (D) RAW264 cells transfected with the indicated HA-Rab5 mutant vector, HA-Rab5 wild type vector or control vector. The cells were incubated with 100 ng/ml LPS for 18 h and then fixed with paraformaldehyde. S-nitrosylated proteins were visualized using a biotin-switch assay (yellow), and HA-Rab5 was visualized after immunostaining with an anti-HA antibody (blue). Scale bar: 5 µm. Fluorescence intensities of nitrosylated protein (yellow) and HA-Rab5 (blue) were measured along the lines in the expanded panel.

### Cysteine residues 212 and 213 in Rab5 are important for Rab5 activity and phagocytic activity

A biotin switch assay was performed using the GST-fused Rab5 mutant to confirm the S-nitrosylation site of Rab5 mediated by NO. Low levels of Rab5 S-nitrosylation were detectable by western blotting for HA-Rab5C212A and HA-Rab5C213A mutant proteins following GSNO treatment, but S-nitrosylation was completely absent for the HA-Rab5C212A/C213A double mutant (Fig. 7C). Through the biotin-switch assay, S-nitrosylated proteins can be biotinylated, allowing their detection not only by Western blotting but also by fluorescence microscopy(82). We also observed S-nitrosylation of Rab5 in a biotin-switch assay using confocal fluorescence microscopy. RAW264 cells transfected with each of the single HA-Rab5 Cys mutants were incubated with LPS for 18h, and then the subcellular localization of S-nitrosylated Rab5 was assessed by a biotin-switch assay and immunostaining. Fluorescence microscopy showed that the HA-Rab5C212A and HA-Rab5C213A single mutants, as well as the HA-Rab5C212A/C213A double mutant, exhibited reduced S-nitrosylation compared to wild type Rab5, which is similar to the western blotting results with the biotin switch assay (Fig. 7D). These results indicate that the two C-terminal cysteine residues, Cys 212 and Cys213, of Rab5 are crucial for S-nitrosylation.

To clarify whether activation of Rab5 by NO involves S-nitrosylation of Rab5, HEK293 cells transfected with the HA-Rab5 C212A and C213A single mutant vectors, HA-Rab5C212A/C213 vector, HA-Rab5 wild-type vector or control vector were incubated with GSNO for 1 h and Rab5 activity was measured in a GST-R5BD pulldown assay. Compared to wild type HA-Rab5, HA-Rab5C212A, HA-Rab5C213A, and HA-Rab5C212A/C213A mutants all had lower Rab5 activity (Fig. 8A).

**Fig 8.**
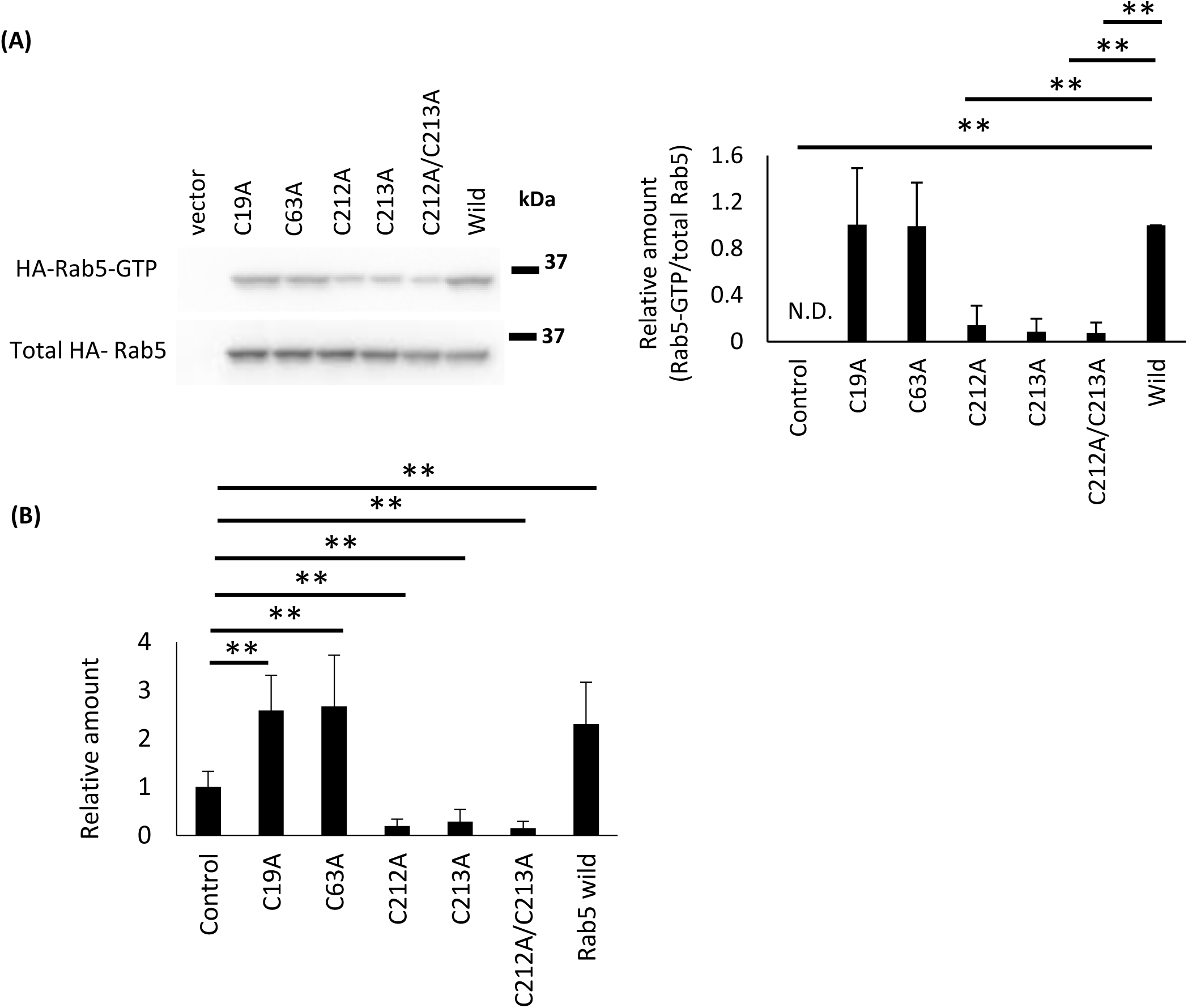
Rab5 cysteine residues C212 and C213 are important for activity and phagocytic activity in RAW264 cells. (A) HEK293 cells transfected with the indicated HA-Rab5 mutant vector, HA-Rab5 wild type vector or control vector were incubated with GSNO for 1h. Rab5 activity was then assessed with a GST-R5BD pulldown assay. Each value in the graph is the mean ± SD of four independent experiments. **p < 0.01 versus Rab5 wild type, N.D., not detected. (B) RAW264 cells transfected with the indicated HA-Rab5 mutant vector, HA-Rab5 wild type vector or control vector were incubated with pHrodo Red *S. aureus* BioParticle conjugates for phagocytosis for 1 h. Phagocytosis of *S. aureus* was measured using a fluorescence microplate reader. Each value in the graph is the mean ± SD of three independent experiment. **p < 0.01 versus vector (mock)-transfected cells.

Next, we examined the effects of each HA-Rab5 mutant on phagocytosis to establish a relationship between Rab5 S-nitrosylation and phagocytosis. Overexpression of HA-Rab5C212A, HA-Rab5C213A, and HA-Rab5C212A/C213A all resulted in markedly decreased phagocytosis activity (Fig. 8B). Taken together, these results indicate that the C-terminal cysteine residues C212 and C213 in Rab5 are important for Rab5 activity and phagocytosis.

### Effect of NO on Rab5 and *S. aureus* clearance in vivo

We further analyzed Rab5 activity and Rab5 S-nitrosylation using LPS, a potent inducer of iNOS, resulting in high levels of NO production in vivo. Mice received tail vein injections with 4 µg/g weight LPS or 4 µg/g weight + 0.1mg/g weight L-NAME. The mice were sacrificed 24 hours later and the spleens were collected. Cytoplasmic fractions of the spleen were prepared and analyzed for Rab5 activity by a GST-R5BD pulldown assay and Rab5 S-nitrosylation was assessed with a biotin switch assay.

The GST-R5BD pulldown assay showed increased Rab5 activity in mice treated with LPS alone compared to untreated mice, while Rab5 activity was increased in mice treated with LPS and L-NAME compared to untreated mice (Fig. 9A). Biotin switch assays indicated enhanced Rab5 S-nitrosylation in mice treated with LPS alone compared to untreated mice, and attenuated S-nitrosylation of Rab5 in mice treated with LPS and L-NAME compared to untreated mice (Fig. 9B). Mouse peritoneal macrophages were also treated with L-NAME for 24 h and then the treated macrophages (5×10^6^) were injected into 8-week-old BALB/c mice together with 3×10^9^ CFU of *S. aureus* ATCC25923. After 6 h, *S. aureus* were collected by peritoneal lavage and plated on tryptic soy agar. The colonies were enumerated as CFUs. Determination of CFUs showed that the number of viable bacteria in untreated cells was very low (6.4×10 CFUs), whereas the number of viable bacteria in the L-NAME-treated cells was very high (6.2×10^3^ CFUs) (Fig 9C). Considering the in vivo experimental results along with the results from cultured cells and in vitro experiments (Fig. 1-8), it is comprehensively likely come up to the conclusion that Rab5 regulation through NO could play an important role in phagocytosis in vivo as well. Based on the findings of this study, a model of the role of S-nitrosylated Rab5 in phagocytosis is shown in Fig. 10.

**Fig. 9.**
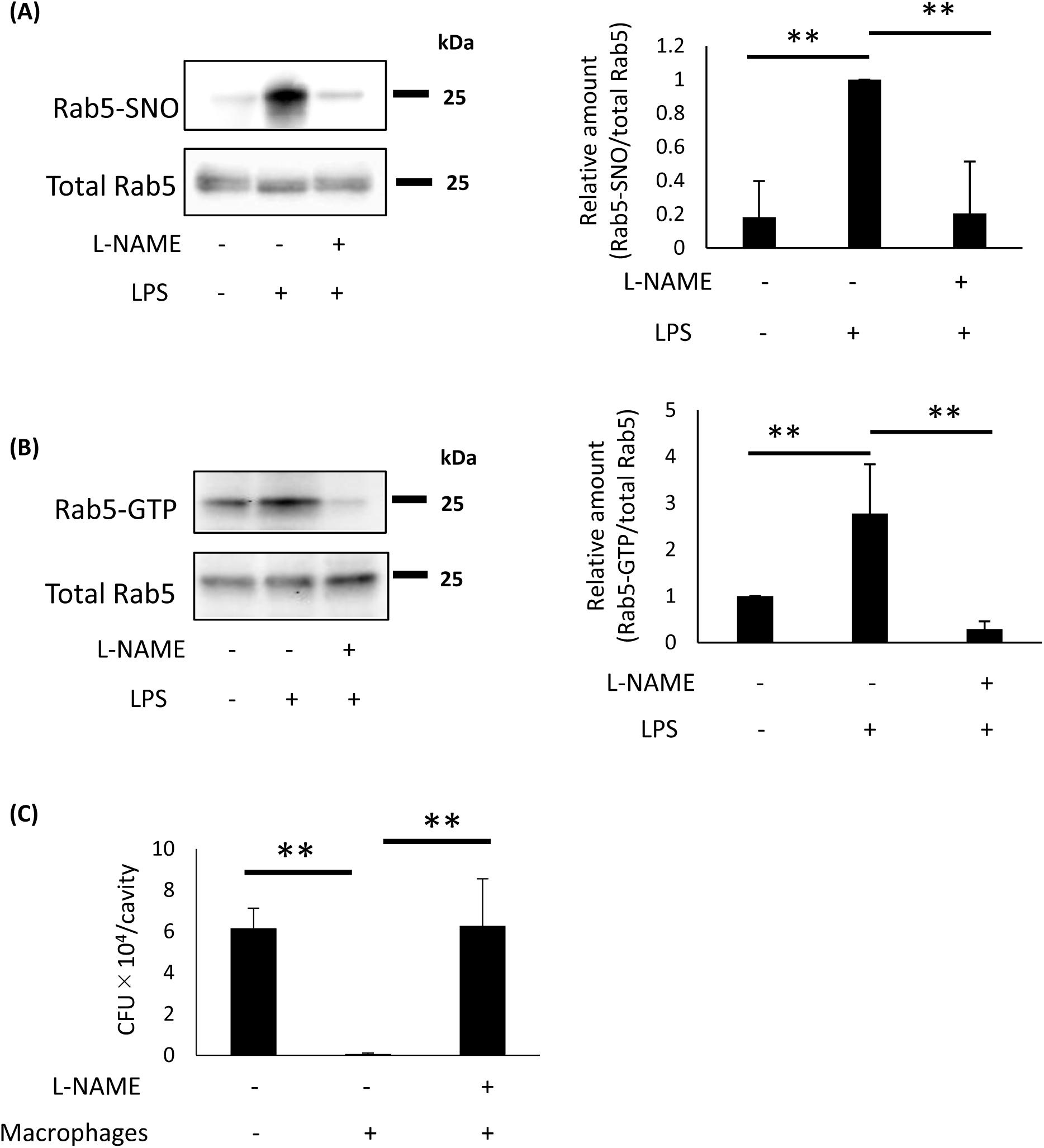
Effect of NO on Rab5 and phagocytosis in vivo. Mice were injected intravenously with LPS or LPS and L-NAME and sacrificed 24 hours later. The spleens were removed and homogenized. The homogenates were analyzed for (A) Rab5 S-nitrosylation using a biotin switch assay and (B) Rab5 activity by GST-R5BD pulldown. Values in the graphs are the mean ± SD of four independent experiments. **p < 0.01. (C) Murine peritoneal macrophages were pretreated with L-NAME for 24 h. Eight-week-old BALB/c mice were injected intraperitoneally with 5×10^6^ L-NAME-treated murine peritoneal macrophages and 3×10^9^ CFU of *S. aureus* ATCC25923. After 6 h, *S. aureus* bacteria were collected by peritoneal lavage and plated on tryptic soy agar. CFU values are the mean ± SD for three or four mice. **p < 0.01.

**Fig. 10.**
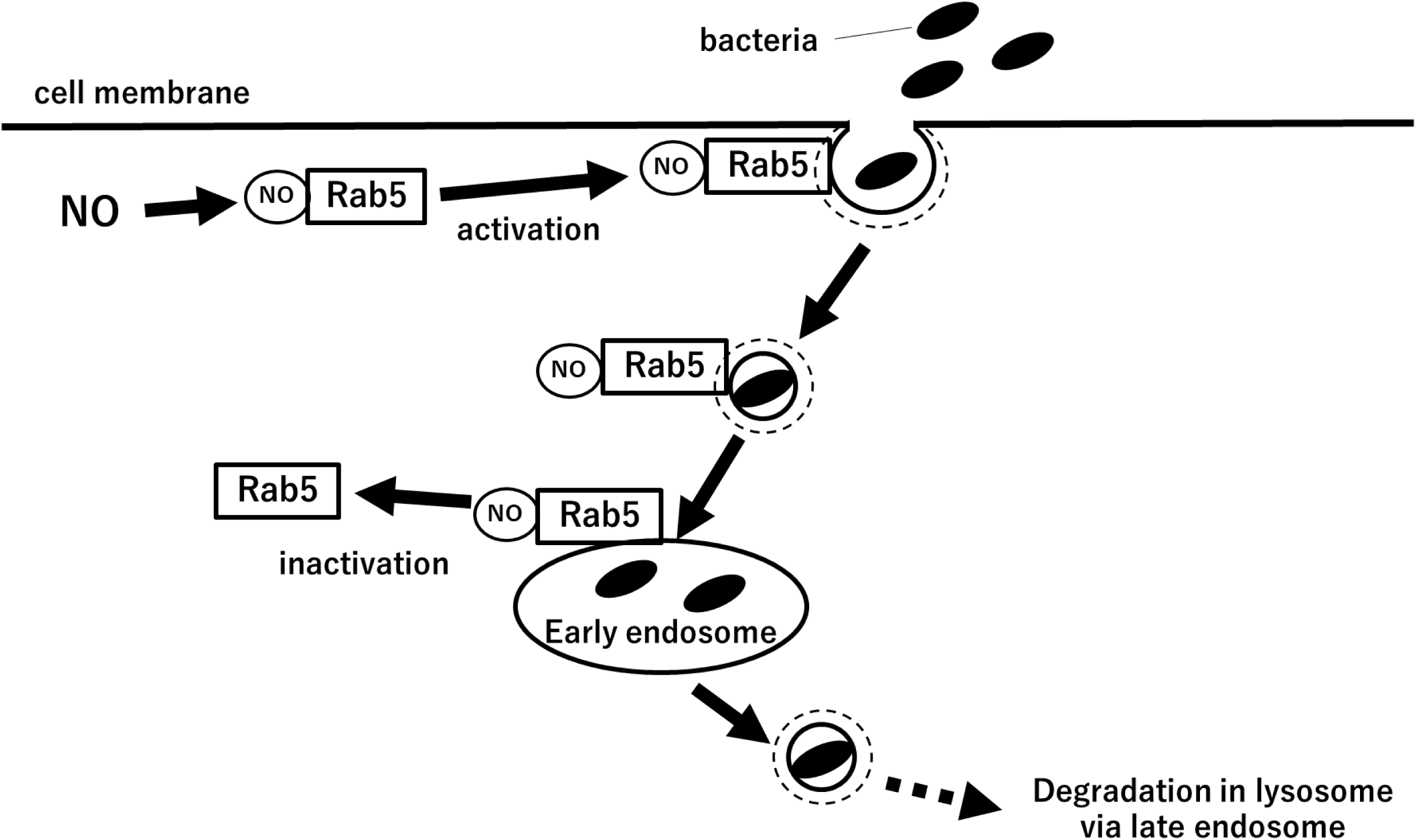
Model showing the role of Rab5 in S-nitrosylation mediated phagocytosis. NO S-nitrosylates Rab5, activating it and promoting phagocytosis. Internalized materials such as bacteria are transported to early endosomes by S-nitrosylated Rab5. Taking other studies together, bacteria would be transported from early endosomes via late endosomes to lysosomes by Rab7. The bacteria are then degraded in the lysosomes.

## Discussion

In the present study, we demonstrate that NO directly activates Rab5 and enhances phagocytosis. Previous reports have shown that GEF activates Rab5, but to our knowledge, this is the first study to show that NO activates Rab5 in the absence of traditional GEF. Moreover, our data newly uncover that phagocytosis is activated by NO, indicating that NO may regulate phagocytosis independently of geranylgeranylation by S-nitrosylating Rab5.

Rab5 is reported to play a role both in clathrin-dependent endocytosis and phagocytosis. In the present study, NO activated Rab5 and promoted phagocytosis (Fig. 1A-F). Within cells, levels of NO produced by iNOS are known to be markedly elevated during bacterial infection(83). NO is known to have direct bactericidal effects, but at levels that are substantially higher than those present under physiological conditions(49). In particular, NO concentrations decrease with increasing distance from iNOS(49). Instead, phagocytosis, in which bacteria are taken up by endocytosis and degraded, is a more effective means to eliminate bacteria during bacterial infection. Previous reports showed that NO activates phagocytes that in turn promote phagocytosis(84, 85), although the detailed molecular mechanisms of this activation are largely unclear. Our results are consistent with previous reports showing that NO activates phagocytosis in phagocytic cells, and we further demonstrated that NO activates Rab5 in phagocytosis.

Protein S-nitrosylation is related to expression levels of eNOS, nNOS, and iNOS in tissues(86, 87) and is restricted to regions of the cell where NOS is localized(58, 59). The production of NO from arginine mediated by iNOS activates macrophages(84, 85), which are essential for host defenses against many pathogens. During phagocytosis, iNOS can be recruited to phagosomes containing *Escherichia coli*(88), *Listeria monocytogenes*(89), or latex beads in IFN- and LPS-stimulated macrophages(90, 91). Our results show that iNOS expression increases during phagocytosis (Fig. 1C and D) and that Rab5 interacts with iNOS (Fig. 2A and B). Moreover, Rab5 is S-nitrosylated in the presence of iNOS expression (Fig. 6D). Based on these observations, Rab5 S-nitrosylation may be enhanced in regions of the cell where iNOS localizes to promote phagocytosis.

NO was previously shown to directly affect NO-mediated GDP/GTP exchange activity of the small GTP-binding protein Ras(74, 75). Dynamin, the founding member of a family of dynamin-like GTPases (DLPs), is implicated in membrane membrane fission and remodeling, and has a critical role in endocytic membrane fission(92). Like Ras, the activity of dynamin-2 is directly regulated by NO through S-nitrosylation(93–95). Here we found that Rab5 is also S-nitrosylated and its GDP/GTP exchange is directly controlled by NO (Fig. 4A and B; Fig. 5A and B). The three-dimensional structures of inactivated (GDP-binding form) and activated small GTP-binding proteins (GTP-binding form) differ(96). Together, these findings suggest that NO may regulate the function of proteins through S-nitrosylation that promotes structural changes that have a direct activating effect on GTP-binding proteins and other G proteins (e.g., dynamin).

The C terminus of many small GTP binding proteins is prenylated by the attachment of geranylgeranyl or farnesyl groups(96). In Rab5, the C-terminal residues C212 and C213 undergo post-translational geranylgeranyl modification that is thought to be important for plasma membrane targeting of Rab5(44, 97). Here we show that NO activates Rab5 and that the C212 and C213 cysteine residues of Rab5 are S-nitrosylated by NO, suggesting that Rab5 may bind to membranes in a prenylation-independent manner (Fig. 7C). Rab13, another Rab family member, is reported to undergo geranylgeranylation of a C-terminal cysteine residue(98). However, Ioannou et al. reported that Rab13 lacking the C-terminal geranylgeranyl binding site still binds to membranes, and proposed a model in which Rab13 binds to membranes via protein-protein interactions(99). Meanwhile, the C terminus of Ras undergoes farnesylation that is thought to be important for membrane binding(100, 101). Ras is also reported to be activated and membrane-bound in a prenylation-independent manner(102). Thus, prenylation is important for membrane binding of small GTP-binding proteins, but this membrane binding can also occur in a prenylation-independent manner. Like Rab13, Rab5 with C-terminal S-nitrosylation may also bind to membranes through protein-protein interactions. Alternatively, multiple small GTPases including Rab5(29, 30, 103), Rab3(104), Rap1(103, 105), and Ras(103) are reported to localize and bind to lipid rafts (detergent-resistant membrane fraction) that are enriched in cholesterol and glycophospholipids(106). A large number of proteins have been found in the detergent-insoluble lipid raft fraction and are thought to bind strongly to lipid rafts through protein-lipid interactions(106). The mechanism by which Rab5 binds to membranes in a geranylgeranylation-independent manner is a subject for future study.

The P fraction of HA-Rab5C212A/C213A mutant showed only one band in the Western blotting results, while two bands were detected in wild-type Rab5 and other mutants (Fig. 7B). In previous reports, it has been shown that prenylated small GTP binding proteins exhibit slower migration on SDS-PAGE compared to their non-prenylated small GTP binding proteins(102, 107). That is so say, in Western blotting, HA-Rab5C212A/C213A mutant would have detected one band of protein that was not prenylated at all, while wild-type Rab5 and the other mutants would have detected two bands of prenylated and non-prenylated protein. In the C fraction, on the other hand, two bands were detected for Rab5C212A/C213A, while only one band was detected for wild-type Rab5 and other mutants (Fig. 7B). We do not know the reason for this difference, but in the field of biochemistry, it is well known that data obtained using Western blotting often show band shifts when a protein undergoes certain post-translational modifications. We speculate that post-translational modifications in Rab5C212A/C213A mutant and wild-type Rab5 or other mutants may be different.

In the present study, we found that NO activates phagocytosis, promotes S-nitrosylation of Rab5, and increases Rab5 activity in vivo (Fig. 9A-C). Based on these in vivo results (Fig. 9) and the findings from in vitro studies and cultured cells (Fig. 1-8), it could be considered that NO plays a role in regulating phagocytosis via Rab5, which is part of immune function, within a living organism. Phagocytosis is important for removal of bacteria (traditional phagocytosis)(1, 2) and dead cells from organisms (recently called efferocytosis)(108, 109). In recent years, increased expression of Rab5 in alveolar macrophages has been reported to enhance efferocytosis, suggesting that Rab5 may also be a critical factor in this process(110). Abnormalities in removal of dead cells due to decreased phagocytosis can lead to autoimmune disease and chronic inflammatory disease(108, 111). Phagocytosis is also reported to play a role in eliminating cancer cells and in neurological diseases such as Alzheimer’s disease and (112–115). Thus, S-nitrosylation of Rab5 could be involved in a variety of diseases beyond just infectious diseases.

In conclusion, here we revealed that NO causes S-nitrosylation of Rab5 and plays a crucial role in the regulation of Rab5 function in phagocytosis. Furthermore, the results suggest that S-nitrosylated Rab5 may regulate phagocytosis in a geranylgeranylation-independent manner. To our knowledge, our study is the first to show that Rab proteins are S-nitrosylated in mammalian cells; it is possible that Rab proteins other than Rab5 are also S-nitrosylated. Further clarification of the relationship between NO and Rab proteins will provide new insight into membrane trafficking and various pathological mechanisms involving these small GTPase proteins.

## Abbreviations

DEA: Diethylamine
DEA-NONOate: diethylamine NONOate
eNOS: endothelial nitric oxide synthase
GAP: GTPase-activating protein
GDI: Rab GDP dissociation inhibitor
GEF: guanine nucleotide exchange factor
GSNO: S-Nitrosoglutathione
iNOS: inducible nitric oxide synthase
L-NAME: NG-Nitro-L-arginine methyl ester
LPS: lipopolysaccharide
nNOS: neuronal nitric oxide synthase
NO: nitric oxide
NOS: nitric oxide synthase

## Acknowledgments

The authors thank Guangpu Li (University of Oklahoma) for the GST-R5BD construct, and Yuji Yamamoto (Tokyo University of Agriculture) for the Rab5 construct. We also thank the Brain Research Institute, Niigata University for use of their ultracentrifuge.

## Author contributions

Makoto Hagiwara and Hiroyuki Tada designed the work, performed experiments, analyzed data and drafted the manuscript. Kenji Matsushita conceived the study, summarized the study, and drafted the manuscript.

## Declarations

